# Anemonefish use sialic acid metabolism as Trojan horse to avoid giant sea anemone stinging

**DOI:** 10.1101/2024.04.22.590498

**Authors:** Natacha Roux, Clément Delannoy, Shin-Yi Yu, Saori Miura, Lilian Carlu, Laurence Besseau, Takahiro Nakagawa, Chihiro Sato, Ken Kitajima, Yann Guerardel, Vincent Laudet

## Abstract

Anemonefish association with giant sea anemone is an iconic example of mutualistic symbiosis. Living inside the sea anemone without triggering the firing of highly toxic nematocysts present at the surface of sea anemone tentacles provides a unique shelter to the fish, which in return, by its territorial aggressiveness, protects the sea anemone from predators. The mechanisms by which the fish avoids triggering nematocysts discharge remain elusive. One hypothesis proposes that absence of sialic acids might disable nematocysts discharge. Here, we verified four predictions about the role of sialic acids in anemonefish protection: (i) sialic acid levels are lower in anemonefish mucus than in non-symbiotic and sensitive damselfish mucus; (ii) this decrease is specific to mucus and not observed in other organs; (iii) during post-embryonic development the levels of sialic acids are inversely correlated with the level of protection; (iv) the levels of sialic acids are minimal in sea anemone mucus. Taken together, our results allow us to propose a general model, in which anemonefish specifically regulates the level of sialic acids in their mucus to avoid nematocysts discharge. Our analysis also highlights several genes implicated in sialic acid removal as potential targets for allowing protection. Interestingly, our results also suggest that unrelated juveniles of damselfish (*Dascyllus trimaculatus*) capable to live in proximity with giant sea anemone may use the same mechanisms. Altogether, our data suggest that clownfish use sialic acids as a Trojan horse system to downplay the defenses of the sea anemones and illustrate the convergent tinkering used by fish to allow a mutualistic association with their hosts.

**Significance statement:** The mutualistic relationship between anemonefish and giant sea anemones, where the fish shelter among the anemone’s tentacles while defending it from predators, is a classic example of symbiosis. However, how the fish avoids triggering the anemone’s venomous nematocysts has remained a mystery. This study reveals that the fish decrease the levels of sialic acids in their mucus, potentially preventing nematocyst discharge. This finding shed light on the mechanisms underlying this symbiotic relationship. Moreover, the discovery that unrelated damselfish may employ similar strategies underscores the broader significance of convergent adaptations in facilitating similar mutualistic associations in marine fishes.

## Introduction

Symbiosis, the intimate long-term association between two or more organisms of different species, is a fascinating biological phenomenon. Such an association can be beneficial to both partners (mutualism), to only one of them (commensalism) or even be detrimental to one partner (parasitism) (Dimijan, 2000). One of the most striking examples is the long-term association between anemonefishes and their giant sea anemones hosts (Burke da Silva & Nedosyko, 2016). The 28 species of *Amphiprion* genus, that belong to the Pomacentrids family, have in common the ability to form social groups living in a close association with sea anemones which belong to 3 distinct groups (Burke da Silva & Nedosyko, 2016; Kashimoto et al., 2022, 2024).

Studied since the end of the nineteenth century (Collingwood, 1868), this symbiosis is considered as a mutualistic relationship as the sea anemone provides a protection to the anemonefish thanks to their deadly tentacles whereas, by their aggressive behavior, the anemonefishes repels sea anemone predators. Anemonefish also provide nitrogen and carbon to the host and its endosymbiotic zooxanthellae playing therefore an important role in their nutrition (Cleveland et al., 2011; Cui et al., 2019). This symbiosis has always fascinated scientists for three main reasons. First, the anemonefishes can live safely inside the tentacles of their host known to discharge stinging organelles called nematocysts contained in a cnidocytes (Mebs, 2009). Nematocysts, which are threads releasing a cocktail of neurotoxins once it penetrates its target, are released after a mechanical and/or chemical stimulation occurring at the surface of the cnidocytes (Lotan et al., 1996; Ozacmak et al., 2001; Tardent, 1988). It is clear that anemonefish are influencing somehow the triggering of these events as they succeed in living within the sea anemone tentacles. Second, there is a complex species specificity of this mutualistic relationship since a few anemonefish species live only in one sea anemone species (called specialists, e.g. *Amphiprion frenatus*, *A. sebae*, *A. biaculatus*), whereas other may have between 2 or even 10 possible hosts (called generalists, e.g. *A. ocellaris*, *A. bicinctus*, *A. perideraion*, *A clarkii*) (Fautin & Allen, 1997; Litsios et al., 2014; Kashimoto et al., 2024). Third, this symbiosis is in fact a tripartite association as the giant sea anemone are themselves symbiotic animals that host a symbiotic dinoflagellate algae *Symbiodinium* providing them 80% of their energy via photosynthesis. It has also been shown that the 3 partners are metabolically connected (Hoepner et al., 2022; Verde et al., 2015).

Even though numerous studies have tried to better understand the resistance of anemonefishes to sea anemone stinging, this question remains unresolved (Burke da Silva & Nedosyko, 2016; Hoepner et al., 2022). Three main hypotheses have been proposed: (i) anemonefish have a thicker mucus layer than other fishes that protect them as a shield ; (ii) anemonefish molecularly mimics the composition of anemone mucus and (iii) anemonefish mucus lacks the trigger for firing sea anemone nematocysts. Individual evidence supports each of these claims. For example, it has been shown that *A. clarkii* mucus was three to four times thicker than that of other coral reef fish species, and did not elicit any response from the sea anemone (Lubbock, 1981). Concerning the second hypothesis, it is proposed that anemonefish, by covering themselves with sea anemone mucus would inhibit nematocysts discharge via a similar mechanism used by sea anemone to prevent nematocysts firing on their own tentacles (Elliott et al., 1994). Comparison of anemonefish mucus and sea anemone revealed, for example, the presence of anemone antigens in *A. clarkii* mucus when inhabiting inside its host (Elliott & Mariscal, 1997). It has also been shown that anemonefish and sea anemone microbiome converge after association, providing an argument in favor of this hypothesis and also suggests the potential for microbial proteins to be involved in molecular mimicry (Pratte et al., 2018; Roux et al., 2019). Arguments in favor of the third hypothesis come from genomic analysis that identified genes under positive selection at the base of the anemonefish radiation (Marcionetti et al., 2019). Among these genes, some implicated in sugar biogenesis have been observed, suggesting that difference in mucus composition may have been instrumental for the protection. This is supported by the observation that a sugar, the 5-*N*-acetylneuraminic acid (Neu5Ac) can stimulate cAMP production and activate calcium channels in sea anemone tentacles and have a role in chemo-sensitization of nematocyst discharge (Ozacmak et al., 2001; 2021). In accordance with these observations, it has been shown that *A. ocellaris* mucus is lacking Neu5Ac (Abdullah & Saad, 2015). These data suggests that the lack of Neu5Ac may be part of the explanation of the protection of anemonefish. However, all these scattered data do not allow to have a clear understanding of how anemonefish can live unharmed within the tentacles in contrast to other fishes.

Neu5Ac belongs to a wider class of acidic monosaccharides called sialic acids that are themselves a subset of a family of α-keto acid monosaccharides with a 9-carbon backbone called nonulosonic acids (NulOs). They mostly substitute the non-terminal extremity of numerous glycolipids and protein glycans in all vertebrates. Due to their usual terminal position and their nature, molecules with sialic acid capping are mediators of numerous biological processes such as ligand-receptor and cell-cell interactions (Schauer, 2009). Although Neu5Ac is among the most abundant sialic acid encountered in nature, more than 50 forms of sialic acids have been identified among which 5-*N*-glycolylneuraminic acid (Neu5Gc) and 2-keto-3-deoxy-nononic acid (Kdn) are well represented in most animals (Angata & Varki, 2002; Chen & Varki, 2010; Schauer, 2000). It should be noted that Neu5Ac, Neu5Gc and Kdn have all been identified in numerous species of fishes, although sometimes in an exquisitely organ-specific manner (Aoki et al., 2021; Inoue et al., 1988; Venkatakrishnan et al., 2019; Yamakawa et al., 2018). It is thus highly possible that a precise regulation of the sialic acids present at the fish surface could explain the inability of anemonefish to trigger their host stinging cells.

As sialic acids have been singled out for a possible role in anemonefish protection, we decided to better evaluate their possible importance. If we consider that they have a role in the protection to stinging, we can make four predictions: (i) sialic acid amount in the mucus of several anemonefish species should differ from the mucus of damselfishes that are the most closely related fish and that are triggering sea anemone nematocyst discharge; (ii) as sialic acids are essential for many biological process we would not expect to see their levels affected in other organs; (iii) during anemonefish larval development, we should observed a shift in the level of sensitivity towards sea anemone stinging associated with a shift in sialic acid levels. Indeed, young larvae have been suspected to trigger nematocysts discharge, they should contain more sialic acid than juveniles or adults; (iv) if anemonefish are using a Trojan horse strategy, that is, if the absence of sialic acid prevents nematocysts discharge, we could expect low levels of sialic acids in sea anemone mucus. In this study we have tested, and validated these predictions and we therefore propose that the specific absence of sialic acids on fish mucus explain why anemonefish live unharmed their host tentacles.

## Results

### Prediction 1: Mucus sialic acid composition of anemonefishes is different compared to damselfish mucus

Analysis of sialic acid composition has been conducted both in fishes maintained in husbandry (at the marine station of Banyuls-sur-mer) and in wild caught fishes (in French Polynesia and in Okinawa). The objective was to compare sialic acid levels in the mucus of anemonefish and damselfishes that are the fish species the most closely related to anemonefish and unable to live inside sea anemones.

We analyzed the mucus of 5 lab-maintained anemonefishes species without sea anemone (*A. biaculeatus*, *A. clarkii*, *A. frenatus*, *A. ocellaris*, *A. percula,* n=3 per species) and juveniles of two damselfish species (*Dascyllus trimaculatus* and the non-symbiotic species *Acanthochromis polyacanthus,* n=3 per species). Of note, we used juveniles of *Dascyllus trimaculatus* which, when tolerated by anemonefish, can live very close to giant sea anemones benefiting from their protection. In contrast, adults of this species are associated to corals. The results revealed that the mucus of symbiotic species contains less sialic acids than the non-symbiotic species (Fig. 1A). More specifically, both anemonefish and *D. trimaculatus* mucus have four time less Neu5Ac than *A. polyacanthus* (p-value=0.057), and the deaminated sialic acid Kdn is not detected in anemonefish and *D. trimaculatus* whereas it is detected in *A. polyacanthus* (Fig. 1A). Neu5Gc was not detected in any of the analysed species.

**Figure 1:**
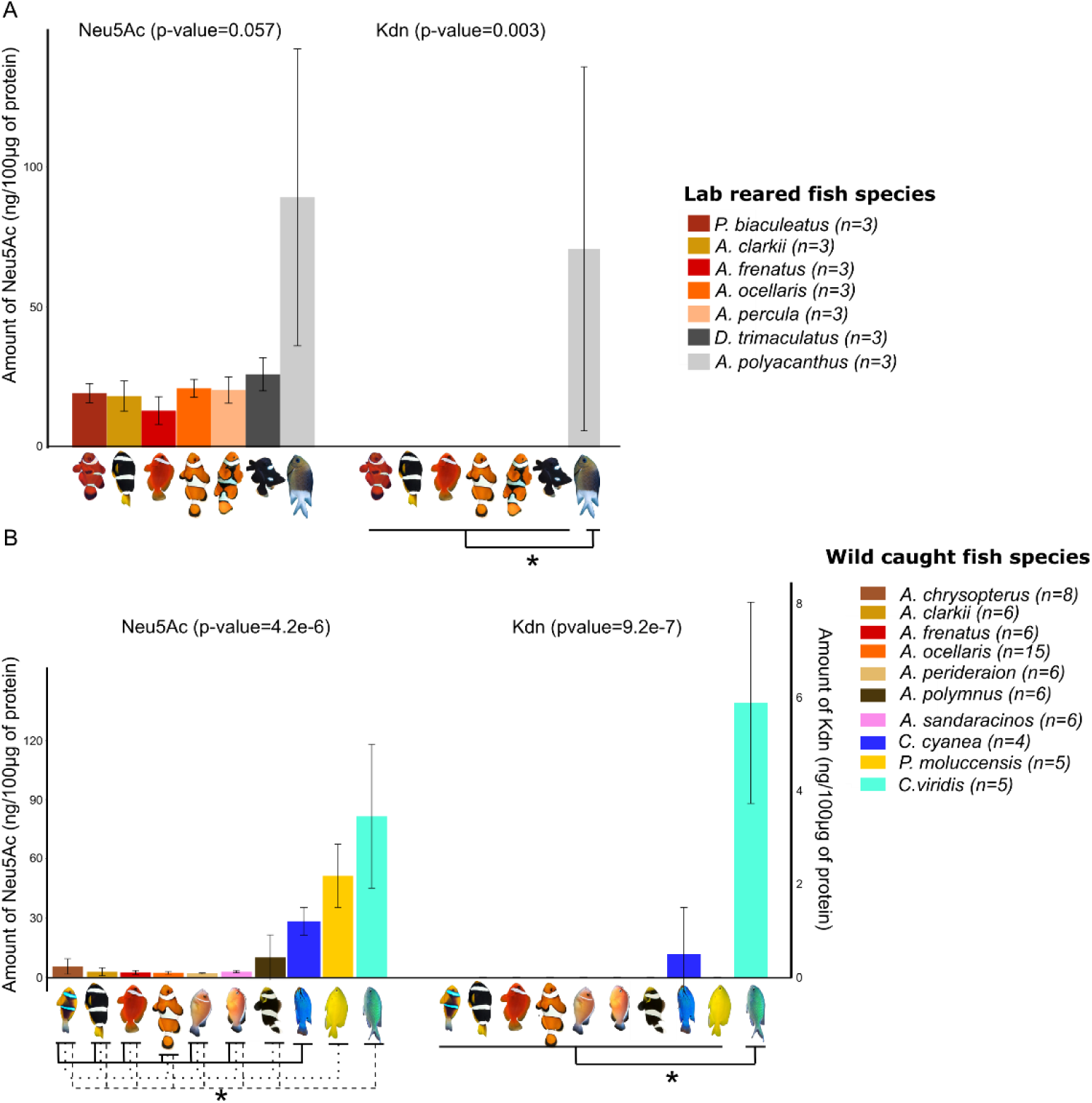
Sialic acid levels in lab and wild caught pomacentridae showing less Neu5Ac in symbiotic species. A) Neu5Ac and Kdn levels (expressed in ng/100µg of protein) in lab reared fish (held without sea anemone): five symbiotic anemonefish species (*Amphiprion biaculeatus, A. clarkii, A. frenatus, A. ocellaris, A. percula*) and 1 symbiotic damselfish species (*Dascyllus trimaculatus*) compared to one non symbiotic damselfish species (*Acanthochromis polyacanthus*). A non-parametric Kruskall-Wallis test (p-value on top of the graph) followed by a pairwise Wilcoxon text were performed. Significant differences between symbiotic species and *A. polyacanthus* are displayed by a star (*). B) Neu5Ac and Kdn levels (expressed in ng/100µg of protein) in wild caught fish: seven symbiotic anemonefish species (*A. chrysopterus, A. clarkii, A. frenatus, A. ocellaris, A. perideraion, A. polymnus, A. sandaracinos*) and three non-symbiotic species (*Chrysiptera cyanea, Pomacentrus moluccensis, Chromis viridis*). A non-parametric Kruskall-Wallis test (p-value on top of the graph) followed by a pairwise Wilcoxon text were performed for both Neu5Ac and Kdn levels. Only significant differences between anemonefishes species and each damselfish species are displayed for Neu5Ac and indicated by a star (*). Only significant difference between *C. viridis* and all the other species are displayed and indicated by a star (*).

As anemonefish maintained in laboratory and used in this study were not associated with sea anemone for practical reasons, we confirmed these results for fish living in natural conditions. We analyzed the sialic acid composition in mucus from fish sampled in the wild, comparing 7 anemonefish species: *A. chrysopterus* from French Polynesia (n=8) as well as *A. clarkii* (n=6), *A. frenatus* (n=6), *A. ocellaris* (n=15), *A. perideraion* (n=6), *A. polymnus* (n=6), *A. sandaracinos* (n=6) and 3 non symbiotic damselfish: *C. cyanea* (n=4), *P. moluccensis* (n=5), *C. viridis* (n=5), all from Okinawa. The results obtained fully confirmed the data observed from lab-maintained fish.

Symbiotic anemonefish species showed significantly less Neu5Ac compared to the three non-symbiotic damselfish species (mean anemonefish Neu5Ac levels comprised between 2.6 and 7.3 ng/100µg of protein whereas mean damselfish Neu5Ac levels were comprised between 28,1 and 81,5 ng/100 µg of protein; p-value<0.05). As previously mentioned for husbandry fishes, no Neu5Gc (data not shown) nor Kdn were detected in anemonefish mucus (Fig. 1B). Neu5Gc was also undetected in the three damselfish, but Kdn was detected in various proportions in *C. viridis* and *C. cyanea* but not in *P. moluccensis* (p-value<0.05, Fig. 1B). Interestingly, when we compared the amount of sialic acid present in fish maintained in laboratory (without se anemone) from those living in the wild (with sea anemone) we observed a statistically significant greater amount in specimens not association with sea anemone (Supp. Fig. 1A).

Taken together these results fully validate the first prediction: sialic acid levels and more specifically Neu5Ac, are always much lower in anemonefish and *D. trimaculatus* (associated with giant sea anemone at juvenile stage) than in non-symbiotic damselfish whatever the conditions or the origins of the fish.

### Prediction 2: Anemonefish display reduced sialic acid content only in mucus compared to other organs

To test the second prediction, we measured sialic acid levels in various organs in comparison to mucus of the symbiotic species *A. ocellaris* (n=10) compared to two non-symbiotics species (*C. cyanea* and *C. viridis*, n=10 per species). Interestingly, the results revealed much higher levels of Neu5Ac in all anemonefish organs compared to the mucus, suggesting that the observed reduction is specific to mucus (Fig. 2A, p-value<0.05). When compared to damselfish organs, *A. ocellaris* Neu5Ac levels were not necessarily lower in all organs. For example, Neu5Ac was significantly higher in *A. ocellaris* skin, muscle, liver and digestive tract compared to *C. cyanea* (Fig. 2B, p-value<0.05). On the contrary, *C. viridis* Neu5Ac levels were significantly higher than both *A. ocellaris* and *C. cyanea* in liver and brain (Fig. 2B, p-value<0.05). Kdn was only detected in *C. viridis* mucus, liver and digestive tract, but not in any organs of *A. ocellaris* or *C. cyanea* (Supp. Fig. 2B).

**Figure 2:**
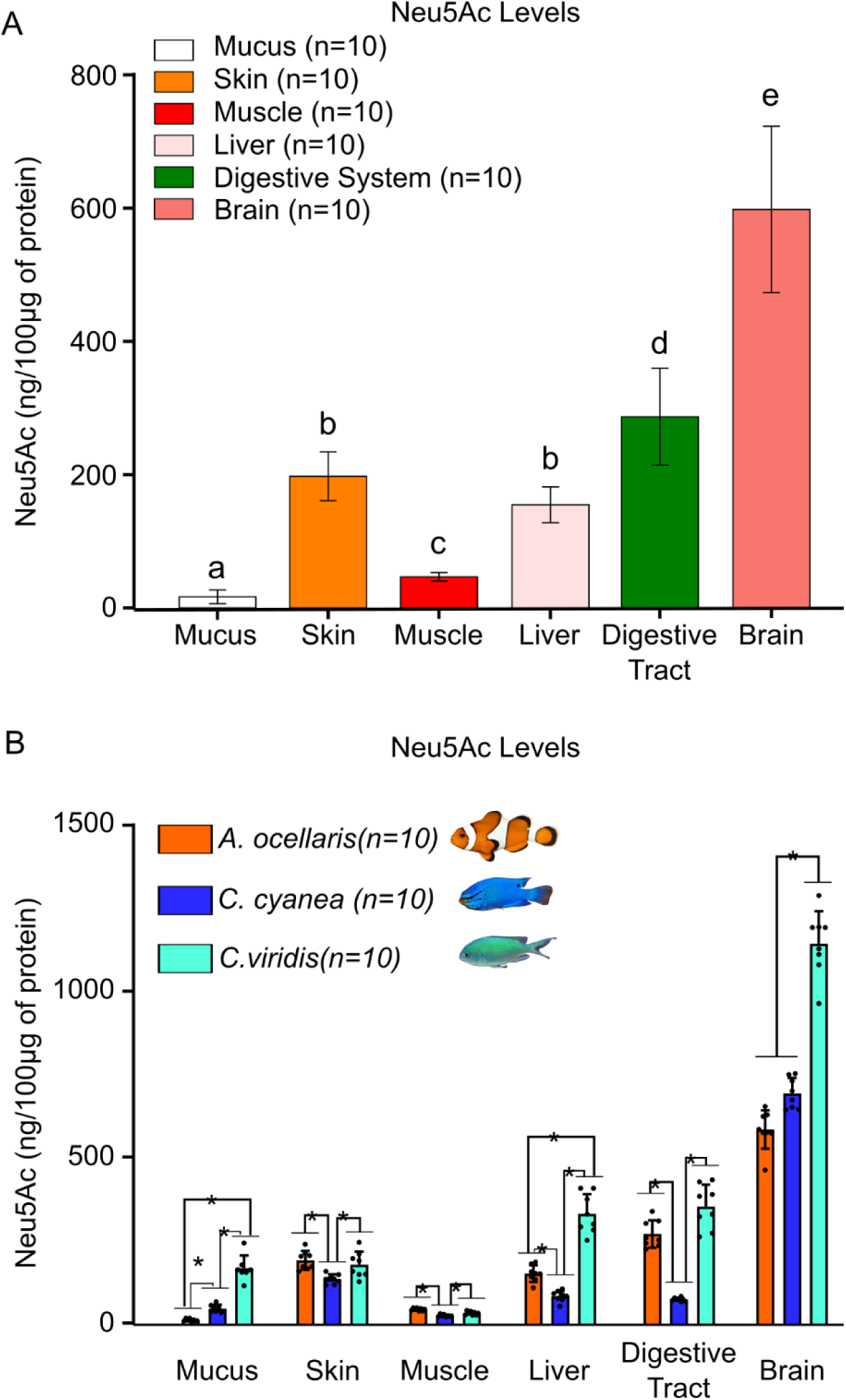
SA levels in pomacentrids various organs confirm a decrease specific to *A. ocellaris* mucus. A) Neu5Ac levels in *A. ocellaris* organs (mucus, skin, muscle, liver, digestive tract and brain). A non-parametric Kruskall-Wallis test (p-value<0.05) followed by a pairwise Wilcoxon test were performed to compare Neu5Ac levels between *A. ocellaris* organs. Organs displaying different letter are significantly different. B) Neu5Ac levels in *A. ocellaris*, *C. Cyanea*, *C. viridis* organs (mucus, skin, muscle, liver, digestive tract and brain). Depending on the organs a one-way ANOVA or a non-parametric Kruskall-Wallis test was performed followed by a Tuckey HSD or a pairwise Wilcoxon test for each organ to compare Neu5Ac levels between the three species. Significant differences are displayed by a star (*).

These results support the second prediction: sialic acid levels are specifically decreased in anemonefishes mucus but not in other organs, including skin, which demonstrates that the expression of sialic acids is exquisitely regulated in tissues and organs to fulfill their functions. The fact that skin sialic acid level is not different between anemonefish and *C. viridis* has interesting implications that will be discussed later.

### Prediction 3: Resistance towards stinging is acquired during metamorphosis and correlates with sialic acid content

Although it has always been claimed that young anemonefish larvae are sensitive to sea anemone tentacles, no clear evidence has been brought for confirmation. To unequivocally determine when clownfish starts to become resistant towards sea anemone stinging, larvae were sampled at each of the 7 developmental stages (previously described in Roux et al., 2019), put in contact with the giant sea anemone *S. gigantea* and survival rates were recorded for each stage. We observed that young larvae (Stage 1 and 2) are highly sensitive towards stinging as non-survived after contact (Fig. 3A). Survival rates started to increase at stage 3 and 4 (10%, 50% respectively, Fig. 3A) and reached 100% at stage 6 and 7. Stage 4 marks the onset of metamorphosis and the transition between the oceanic dispersal phase and the reef phase (Roux et al., 2019, 2023). After entering in a reef, anemonefish larvae must locate a reef and then a suitable sea anemone to settle. It is thus necessary for them to be able to enter their host without being stung which is what has been observed here. Once larvae start metamorphosing, meaning when they are ready to settle into a sea anemone host, survival rates increased.

**Figure 3:**
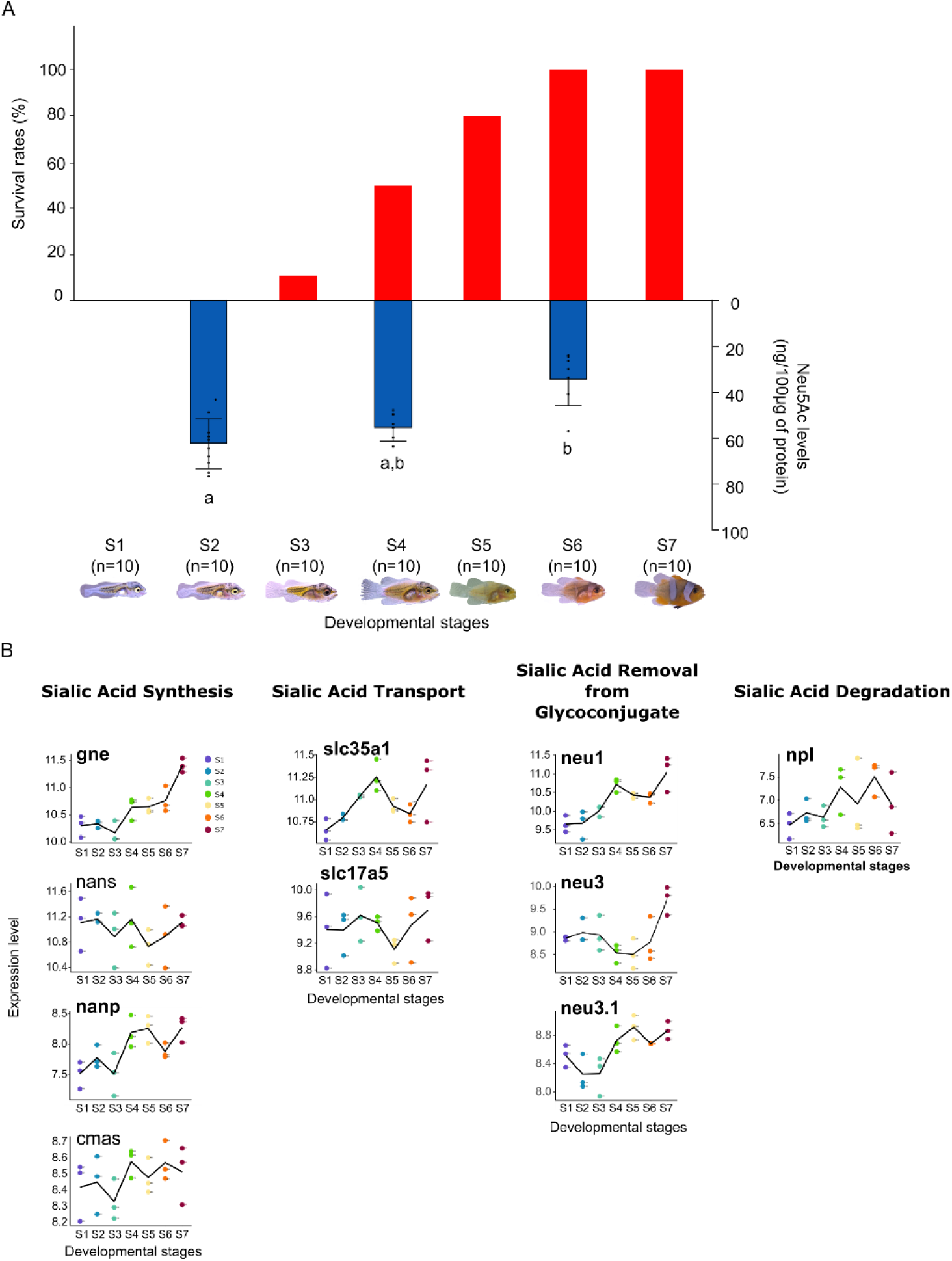
Survival rates and sialic acid production changes during *A. ocellaris* larval development. A) Combined graphs showing in red the survival rates (percentage) of *A. ocellaris* larvae sampled at different developmental stages (Roux et al., 2023) and in blue Neu5Ac levels (expressed in ng/100µg of protein) of larvae sampled before metamorphosis (stage 2) and during metamorphosis (stage 4 and 6). A one-way ANOVA followed by a Tuckey HSD test was performed on Neu5Ac levels to compare each developmental stage. Stage with different letter have significantly different Neu5Ac levels. B) Expression levels of genes involved in Sialic acid synthesis (*gne, nans, nanp, cmas*), transport (*slc35a1, slc17a5*), removal from glycoconjugate (*neu1, neu3, neu3.1*), degradation (*npl*). Genes written in bold are significantly differentially expressed between pre metamorphosis stage (S1 and/or S2 and/or S3) and metamorphosis stages (S5 and/or S6 and/or S7).

In addition to assessing survival rates, we also measured the levels of sialic acid on stage 2, 4 and 6 larvae to determine if the survival rates increase might be corroborated with a change in the sialic acid composition of the larvae. On the three sialic acids commonly measured (Neu5Ac, Neu5Gc and Kdn) only Neu5Ac was detected, and we observed a significant decrease at stage 6 compared to Stage 2 and 4 (*p-value* <0.05, Fig. 3B). It is worth to note that, because of the size of clownfish larvae, mucus collection was not possible, therefore we compared the sialic acid content in the whole animal. Despite this experimental limitation these results could be interpreted as a confirmation of the third prediction.

These data suggest that resistance is acquired during metamorphosis. As the transcriptomic changes occurring during anemonefish larval development and metamorphosis have been previously analyzed (Roux et al., 2023), we scrutinized the expression of sialic acid metabolism genes during metamorphosis to eventually extend the correlation between protection and sialic acids. For this, transcriptomic data obtained by Roux et al., 2023 for each developmental stages (entire larvae) were used to retrieve the normalized expression levels of 33 genes involved in Neu5Ac metabolism. These genes were categorized into 5 genes sets: Neu5Ac synthesis, Neu5Ac transport, Neu5Ac degradation, Neu5Ac removal from glycans as well as Neu5Ac transfer and fixation on glycans. The complete pathway and the expression pattern of each gene are presented in Supp. Fig. 2 & 3. We observed quite heterogenous change of expression, which is not surprising since, because of the small size of the larvae, these transcriptomic data were obtained from entire fish in which sialic acid have important roles in various organs (as shown in Fig. 2). It is therefore not surprising to see such tight regulation of key metabolic genes during larval development. However, as exemplified in Fig 3B, an increased expression in early juvenile stages is observed for 3 genes encoding for enzymes implicated in sialic acid degradation called neuraminidase (namely *neu1, neu3* and *neu 3.1*, Fig. 3B). The gene encoding for neuraminidase 1 (*neu1*) peaks at stage 4 and later increases in juveniles is particularly interesting as it cleaves sialic acid from substrate such as glycoproteins and could therefore be used to remove sialic acid from proteins expressed by mucous cells. The 2 genes *neu3* and *neu3.1* have similar expression patterns, with strong increases until stage 7and could also be implicated in similar activities. This may indicate that the removal of sialic acid is an active phenomenon that is established during metamorphosis, ensuring that the juvenile emanating from metamorphosis can enter safely in a sea anemone (see Discussion).

### Prediction 4: Low level of sialic acids is sea anemone mucus

The fourth prediction is that anemonefish may in fact use the very same mechanism that the sea anemone are using to avoid nematocyst discharge against their own tentacles. Nematocysts discharge, as well as toxins synthesis, has a metabolic cost for sea anemone (Fautin 2009, Sachkova et al. 2020, Kaposi et al. 2022) and it is likely that these organisms have developed a system to not stung themselves. One relevant possibility would be that giant sea anemone also use the lack of sialic acid to avoid stinging themselves and in that view, anemonefish may in fact behave as Trojan horse highjacking the very same strategy to avoid triggering nematocyst discharge. This would imply that giant sea anemone would also have low levels of sialic acid. For this reason, we compared sialic acid levels in four giant sea anemones (*Heteractis magnifica*, *Stichodactyla gigantea*, *Heteractis crispa* and *Entacmaea quadricolor*) that represents the three main types of giant sea anemone associated to anemonefish (Kashimoto et al., 2022).

We did not observe significant amounts of NeuAc, NeuGc, Kdn or any other type of sialic acid in the mucus sampled of these four species (Supp. Fig. 4). Some signals corresponding to potential sialic acids were detected but their intensity was below the detection threshold. These data therefore fully confirm the fourth prediction, suggesting that the absence of sialic acid in sea anemone mucus may be linked to their protection against their own nematocysts and that anemonefish use this system as a Trojan horse to avoid being stung.

## Discussion

Known since the 19^th^ century (Collingwood, 1868), the symbiotic relationship between giant sea anemone and anemonefish still holds secrets for the scientific community who is eager to understand how these fish are able to live unharmed in their deadly host. Our study brings new elements to this question and reinforce the hypothesis that the lack of sialic acids, and more specifically Neu5Ac, avoid anemonefish to trigger nematocysts discharge of their host.

### Sialic acid levels negatively correlate anemonefish sensitivity towards sea anemone stinging

A novel finding of our study is that the ability of anemonefish to live safely in the sea anemone tentacles is acquired during the larval development. As most of the marine fish, anemonefish life cycle is composed of an oceanic larval dispersal followed by a sedentary coastal phase. They reproduce in the vicinity of their sea anemone host, laying eggs on a substrate close to the sea anemone. Right after hatching, larvae are transported into the open ocean and are not supposed to enter in contact with the sea anemone. However, to the best of our knowledge no studies have directly investigated if anemonefish larvae were innately resistant to their host stinging. Interestingly, we demonstrated that the decrease of Neu5Ac levels is negatively correlated with the increase of survival rates of *A. ocellaris* larvae: Our results indeed clearly showed that newly hatched larvae of *A. ocellaris* are highly sensitive to sea anemone stinging as survival rates are low at the beginning of the development. However, survival rates reached 100% at the end of the larval development. The fact that Neu5Ac level followed a reverse tendency compared to survival rates and decreased during larval development clearly indicated that sensitivity towards sea anemone stinging might be linked to Neu5Ac levels. Additionnaly, as sialic acid levels differ in lab-maintained anemonefish when compared to those associated with anemone also suggest that acclimation processes, implicating sialic acid, may also play an important role in the protection mechanisms.

### Anemonefish specifically decrease sialic acid levels in their mucus

By comparing the levels of sialic acids between 9 species of anemonefish (*A. biaculeatus, A. clarkii, A. frenatus, A. ocellaris, A. percula, A. chrysopterus, A. perideraion, A. polymnus, A. sandaracinos*), one species of symbiotic juvenile damselfish (*Dascyllus trimaculatus*), as well as 4 non symbiotic damselfish (*Acanthochromis polyacanthus, Chrysiptera cyanea, Pomacentrus moluccensis, Chromis viridis*) sampled either in a lab environment or in the wild, we also corroborated and extend the results obtained by Abdullah et al., (2015): anemonefish have less sialic acids than non-symbiotic damselfish species. In addition, in accordance with the fact that sialic acids have important biological functions (Schauer, 2009) we observe that this decreased level is tissue specific and affect only the mucus of anemonefish.

### Anemonefish use sialic acid as a Trojan horse to get protection from sea anemone tentacles

The analysis of the sialic acid composition of the mucus isolated from sea anemones did not allow us to observe significant amounts of NeuAc, NeuGc, Kdn or any other type of sialic acid (Supp. Fig. 4). Although some signals corresponding to potential sialic acids were observed, their intensity was below the detection threshold that would allow them to be distinguished from background noise. These experimental results are not in accordance with what has been obtained by Abdullah et al., (2015) which detected high amount of sialic acids. However, the analysis method used in this study, based on thiobarbituric acid, is not specific to sialic acids and is known to be reactive with other compounds like RNA and oxidized lipids. On the contrary, as the method used in our study is highly specific to sialic acids detection, we are confident that the data obtained on sea anemones are in accordance with the fact that sialic acids have never been so far conclusively detected in another species of cniderians (from our knowledge). Further investigation should be carried on investigating the presence of genes involved in sialic acid metabolism in cnidaria to validate our results.

Nematocysts discharge (and toxins synthesis) has a metabolic cost for sea anemone (Fautin 2009, Sachkova et al. 2020, Kaposi et al. 2022). It is thus very likely that these organisms have developed a system to not stung themselves as proposed originally by Schlichter who suggested that anemone mucus contains inhibitory substances that prevent self-stimulation and nematocyst discharge, and that anemonefishes acquire these substances during acclimation (Elliott et al., 1994; Schlichter, 1976). Based on the results obtained in our study, the model would be different in that it suggests that sea anemone have low levels of Neu5Ac in their mucus to avoid auto-stinging and that anemonefish highjacked this system to enter safely into their host.

### What could be the mechanisms at play?

How the anemonefish decreases the amount of sialic acid in their mucus is still unclear and may result from the combination of several mechanisms. First, it has been shown previously that among genes positively selected at the base of the anemonefish radiation and therefore potentially involved in the symbiosis establishment, two encodes for proteins with a functional link with N-acetylated sugars: versican core protein (*vcan*) and the O-GlcNAc transferase (*ogt*) (Marcionetti et al., 2019). Versican core protein is known to be a critical extracellular matrix regulator of immunity and inflammation (Wight et al., 2020) that interacts with several matrix molecules including glycosaminoglycans containing N-acetylhexosamine (Wu et al., 2005). Expression of versican core protein in clownfish skin is thought to bind to N-acetylated sugars and could therefore mask them to the host chemoreceptors therefore preventing nematocyst discharge. Two *vcan* genes were identified in our transcriptomic data and one, *vcana,* who is expressed in the epidermis, showed an increase from stage 4 (marking the onset of metamorphosis, Supp. Fig. 5) suggesting it may play a role in this key period where the young fish acquire the resistance (Marcionetti et al., 2019; Roux et al., 2023). On the other hand, protein O-GlcNAse has the potential to cleave N-acetylated sugars from different cell surface molecules (Bathina, 2014) and has also been found to be expressed in anemonefish epidermis (Marcionetti et al., 2019). One *ogt* gene has been identified in the transcriptomic data set used in this study (*ogt1*) and its expression level decreased from stage 3 in whole fish (Supp. Fig. 5) but we still don’t know how its expression is regulated in skin during metamorphosis.

A second possible mechanism could be the direct tissue specific regulation of genes implicated in sialic acid biochemistry that is sialyltransferases that transfer sialic acid to nascent oligosaccharide, but also neuraminidases (also called sialidase), that remove sialic acids from glycoconjugates. Despite coming from entire individuals and therefore not representative of the exquisite tissue-specific regulation probably at play, the expression levels provide some interesting hints that can be explored in further studies. Indeed, we observed an activation of the pathways governing removal of Neu5Ac exemplified by an increased expression of *neu1*, *neu3* and *neu3.1* coinciding with metamorphosis, the increase of survival rates and the decrease of Neu5Ac levels. Further experiments are required to address gene expression in a tissue specific manner to understand the regulation mechanism involved in the reduction of sialic acid levels in anemonefish.

Another relevant hypothesis that could explain the local regulation of sialic acid content in mucus could be the action of bacteria involved in sialic acid removal. Indeed, several bacteria are known to possess enzymes (sialidases and neuraminidases) involved in sialic acid removal (Li & Chen, 2012) and it has been shown that the mucus of fish and sea anemone converge after contact in terms of microbiota (Pratte et al., 2018; Roux et al., 2019). This is an interesting possibility as it would explain why in many case naive anemonefish need to acclimate to their sea anemone and would be also consistent with the model suggesting a chemical mimicry of anemonefish with sea anemone mucus. An interesting path to follow would be therefore to test if host bacteria are able to remove sialic acids from fish mucus.

### Other possible mechanism at play

Our data suggest that the low level of sialic acid play an important role in the protection mechanism that allow anemonefish to live unharmed inside sea anemone tentacles. However, we do not claim that this lack of sialic acid is the only mechanism at play. This does not explain why some anemonefishes need to acclimate during several minutes before entering in a sea anemone, suggesting that a second process, likely the chemical mimicry, is at play. Recent results suggest that the sea anemone, that have a direct interest to host anemonefish because of territorial defense and metabolic exchange, may also play a role in allowing anemonefish to settle by discharging fewer nematocysts at familiar anemonefish after delayed mucus adaptation (Hoepner et al., 2024).

It is also known that nematocysts can fire following chemical stimulation (like the presence of sialic acid) but also via the activation of mechanoreceptors, it is thus very likely that other unknown mechanism also participates to the protection. For example, there may be difference in the organization of the epidermis between anemonefish and damselfish such as epidermis thickness, scale organization and thickness, amount of mucus etc. that allow the anemonefish to be better protected against low nematocysts discharge that would normally harm or even kill another fish. In other word, it is likely that the lack of sialic acid in the mucus is necessary but is coupled with additional mechanism.

Another important aspect that is not explain by our results is the specificity of association between the 28 species of anemonefish and the ca. 10 species of giant sea anemone. It is clear that there are complex and still largely unknown rules of association with some species of anemonefish being specialists some other being generalist (Hoepner et al., 2022). In our study, we also did not note any difference in sialic acid levels when considering anemonefish host specificity. For example, the generalist *A. clarkii* showed similar levels of Neu5Ac when compared to the specialist species *A. frenatus* (2.69 ± 2.03 (sd) ng/100µg of protein, 2.31 ± 0.98 (sd) ng/100µg). Therefore the model we tested in this study does not explain these effects that are likely due in part to ecological preference, chemoattraction mechanisms but that could also be linked to a sensitivity of some species to the toxic compounds released by the sea anemone (Nedosyko et al., 2014; reviewed in Hoepner et al., 2023).

### Is this a general phenomenon?

One intriguing result of our study is the fact that the juvenile of domino damselfish (*Dascyllus trimaculatus*) also contains low levels of sialic acid in their mucus (Fig. 1A). This species is known to live associated with sea anemone when accepted by dominant anemonefish (Hayashi et al., 2020). However, it is not closely related to anemonefish (Tang et al., 2021) which clearly suggest a case of convergence and reinforce the association between the lack of sialic acid and the protection.

In this context, it is interesting to note that several species of fish, such as cardinalfishes (Apogonidae), wrasses (Labridae), hawkfishes (Cirrhitidae), butterflyfishes (Chaetodontidae), a scaled blenny (Clinidae), and even a temperate greenling (Hexagrammidae), also live associated with sea anemones (Feeney et al., 2019; Karplus, 2014; Randall & Fautin, 2002). Many of those are in fact not sensititive to nematocysts, and can have lesions after contact with the tentacles. In other cases such as the labrisomid *Starksia hassi* or the cardinal fish *Apogon moluccensis,* no apparent lesions are observed despite full contact with the tentacles suggesting that once again a mechanism avoiding nematocyst discharge exist. It is also worth to note that many invertebrates such as shrimps or crabs are living permanently inside sea anemone, as anemonefish (Gusmão & Daly, 2010; Mebs, 2009). Another interesting case, while not being a symbiosis, is the case of nudibranch that feed on cnidarians as they must defend themselves from the prey’s nematocysts and it has been shown that their mucus inhibit the discharge of nematocysts from sea anemone tentacles (Greenwood et al., 2004). It will be interesting to study these cases to see if the convergence observed within damselfishes extend to other species.

In conclusion, our study provides compelling evidence supporting the hypothesis that anemonefish employ sialic acids as a protective mechanism against nematocyst discharge from their giant sea anemone hosts. Remarkably, our observations also suggest a shared utilization of these mechanisms by unrelated damselfish juveniles, underscoring the broader ecological significance of convergent adaptations in facilitating mutualistic interactions within marine ecosystems. Overall, our research offers valuable insights into the intricate mechanisms of the relationships between unrelated organisms and opens avenues for further exploration in symbiosis biology.

## Material and Methods

### Mucus sampling

A total of seven species were sampled in Banyuls sur mer Marine station husbandry: five anemonefishes (*A. biaculeatus* n=3, *Amphiprion ocellaris* n=3*, A. percula* n=3*, A. clarkii* n=3*, A. frenatus* n=3), one damselfish inhabiting sea anemone at juvenile stage (*Dascyllus trimaculatus* n=3), and one non symbiotic damselfish (*Acanthochromis polyacanthus*). Each species was maintained in closed recirculatory system filled with artificial sea water (Red Sea salt, Antinéa, France) and without sea anemones. Temperature was maintained at 26°C, salinity at 34g/L and a 14/10 hours light/dark photoperiod was applied.

A total of ten species were sampled in the wild: one species in Moorea French Polynesia (*A. chrysopterus* n=8), five species in Okinawa Island Japan (*A. clarkii* n=6, *A. frenatus* n=6, *A. ocellaris* n=15, *A. perideraion* n=6, *A. polymnus* n=6, *A. sandaracinos* n=6), and three non symbiotic damselfish species, also encountered in Okinawa (*Chrysiptera cyanea* n=4*, Pomacentrus mollucensis* n=5*, Chromis viridis* n=5). Fish were sampled whether by snorkeling or diving using hand nets.

Mucus collection was conducted always using the following protocol with sterile material and gloves to avoid contamination of the samples. Fish were individually anesthetized in MS222 (200mg/L, Sigma aldrich) and transferred in a glass petri dish without water. Sterile cell scraper (SARSTEDT, Nümbrecht Germany) was used to gently scrap each flank of the fish from gills to tail (5 times per side without touching the gills). The fish were then gently placed back in a container filled with sea water for awakening and place back in its aquarium or released in the wild. Mucus was washed off from cell scraper and petri dish with 2ml of ultrapure water, transferred in a glass tube and kept at −20°C until extraction and analysis. Gloves were change between species to avoid any contamination.

### Organ sampling

To determine if sialic acid composition in anemonefish is decreased in the mucus or in the entire body, fish maintained in aquaria were dissected and the following organs were sampled: mucus, skin, muscle, liver, digestive tract and brain. The anemonefish *A. ocellaris* as well as 2 non symbiotic damselfish *C. cyanea*, and *C. viridis* were sampled for comparison (n=10 per species). *A. ocellaris* juveniles were obtained from OIST husbandry and juveniles of both *C. cyanea* and *C. viridis* were obtained from a local petshop (Makeman, Uruma city). Fish were euthanized in MS222 (400mg/L) and transferred in a glass petri dish for mucus collection (described above) and organ dissection. Organs were stored separately in 1.5ml Eppendorf tubes and kept at −20°C until extraction and analysis. Dissection tools and petri dish were rinsed and disinfected with ethanol between each individual to avoid contamination between samples.

### Sialic Acid Hydrolysis and DMB Derivatization

All samples were lyophilized before extraction. Once lyophilized, samples were incubated in CHAPS extraction buffer (8M urea ; 2% CHAPS, 50 mM DTT, 1X protease inhibitor) and maintained under constant agitation at 4°C overnight. Protein extracts were then centrifuged at 20,000 g, 4°C for 10 minutes and supernatants were collected. Protein concentration was determined by the Pierce™ BCA Protein Assay Kit - Reducing Agent Compatible, according to the manufacturer’s instructions. 40 µg of protein extract were loaded into preconditioned HTS 96-well plates with hydrophobic Immobilon-P PVDF membrane and incubated for 30 min at 37 °C. The wells were washed 6 times with 200 µL mQ water prior to centrifugation (1 min, 500 g). Sialic acids attached to glycoconjugates were released at 60°C for 3h in 0.1M trifluoroacetic acid. Released sialic acids were collected by centrifugation (1 min, 1,000 g) and lyophilized. They were then subsequently coupled to 1,2-diamino-4,5-methylenedioxybenzene dihydrochloride (DMB). Samples were heated at 50 °C for 2 h in the dark in 7 mM DMB, 1 M β-mercaptoethanol, 18 mM sodium hydrosulfite in 5 mM acetic acid. Sialic acids coupled to DMB (DMB-Sia) were then analyzed by liquid chromatography fluorescence detector (LC-FLD).

### Quantitation Analysis of DMB-Sia on LC-FLD

DMB-labeled sialic acids were injected into the Prominence LC-20AB micro LC system (Shimadzu). Samples were applied to an analytical LC column (InfinityLab Poroshell 120 EC-C18, 4.6 x 150 mm, 2.7 µm) and separated isocratically by a solvent mixture of acetonitrile/methanol/water (9:7:84) and identified by referring to the elution positions of standard Neu5Ac, Neu5Gc, and Kdn derivatives. Individual sialic acid derivatives were quantified by integration of fluorescence signals after HPLC separation, plotted against standard curves of corresponding authentic standards.

### Survival experiment

To determine at which stage anemonefish larvae become resistant to sea anemone stinging, larvae of *A. ocellaris* were sampled at each developmental stage (7 distinct stages according to the developmental table of Roux et al., 2019) and put individually using a transparent pipette in contact with the tentacles of the sea anemone *Stychodactyla gigantea* (stage 1 to 5: n=10, stage 6: n=11, stage 7: n=9). Individuals sticked to tentacles and unable to escape were counted as dead and individuals able to freely swim in between the tentacles without sticking to them were counted as surviving. Total number of dead individuals was then used to determine the survival rates for each developmental stage. Larvae were raised in a closed system using natural filtered sea water following methods described in Roux et al., (2021).

### Developmental stage sampling

To assess the sialic acid composition and quantity during *A. ocellaris* larval development, three developmental stages were sampled according to the developmental table of Roux et al., (2019): before metamorphosis (stage 2 n=10), beginning of metamorphosis (stage 4 n=10), and during metamorphosis (stage 6 n=10). Samples were frozen and processed as described above.

### Sea anemone mucus sampling

Four giant sea anempones (*Heteractis magnifica, Stichodactyla gigantea, Heteractis crispa and Entacmaea quadricolor*) that represents the three main types of giant sea anemone associated to anemonefish (Kashimoto et al., 2022). Animals were collected from the wild in Okinawa and maintained in a natural sea water open circuit in OIST marine station. Each specimen was gently caught with a bucket filled with sea water from its own tank and gently brought to the surface to emerge tentacles. Mucus was then collected by putting a glass petri below some tentacles and gently scrapped with a cell scraper. Mucus was then collected and handled similarly as fish mucus sample. Collection was repeated 4 times on different tentacles for each species.

### Sialic acid signaling gene pathway expression level

Expression levels of 33 genes involved in SA pathway were retrieved from each *A. ocellaris* larval developmental stages using the transcriptomic data set published in Salis et al., (2021) and Roux et al., (2023). Those genes were categorized into the following 5 sets. Genes encoding for enzymes involved in Neu5Ac synthesis: *gne* (UDP-GlcNAc 2-epimerase/ManNAc kinase), *nans* (Neu5Ac 9-phosphate synthase), *nanp* (Neu5Ac 9-phosphate phosphatase), *cmas* (CMP-Neu5Ac synthetase). Genes encoding for enzymes involved in Neu5Ac transport: *slc35a1* (CMP-Sia transporter), *slc17a5* (sialin). Genes encoding for enzymes involved in Neu5Ac degradation: *npl* (N-acetylneuraminate pyruvate lyase). Genes encoding enzymes involved in the transfer and fixation of Neu5Ac on sialoglycans (glycoproteins, glycolipids, glycoRNA): sialyltransferases. Finally, genes encoding for enzymes involved in SA removal from sialoglycans: *neu1*, *neu3* and *neu3.1* (neuramindase 1, 3 and 3.1).

### Statistical analysis

Sialic acid levels were analysed using Rstudio software (RStudio Team, 2020). One-way anova was performed for Neu5Ac (p-value on top of the graph) followed by a Tuckey HSD test for post-hoc comparison when parametric test could be used or a non-parametric Kruskall-Wallis test followed by a pairwise Wilcoxon text were performed.

## Acknowledgments and funding sources

We thank Stefano Vianello for critical reading of the manuscript. We also thank the aquariological service of Banyuls Marine station and MEEDU unit in OIST marine station to produce anemonefish and their help for this study. We also thank Kina Hayashi, Manon Mercader, Marleen Klahn, Hiroyuki Takamiyagi, Keishu Asada and Mathieu Reynaud for their help in sampling fish in the wild for mucus collection. We would like to also thank Estelle Garenaux who did preliminary analysis on sialic acid detection and measurements. This work was supported by OIST as well as by an internal Shinka grant funding. We are grateful to the PAGes-P3M core facility (US 41 - UAR 2014 – PLBS) for providing the scientific and technical environment conducive to achieving this work. This study was funded by J-GlycoNet Joint Research Program fund in FY2023.

