## Supplemental Figures for "Anemonefish use sialic acid metabolism as Trojan horse to avoid giant sea anemone stinging"

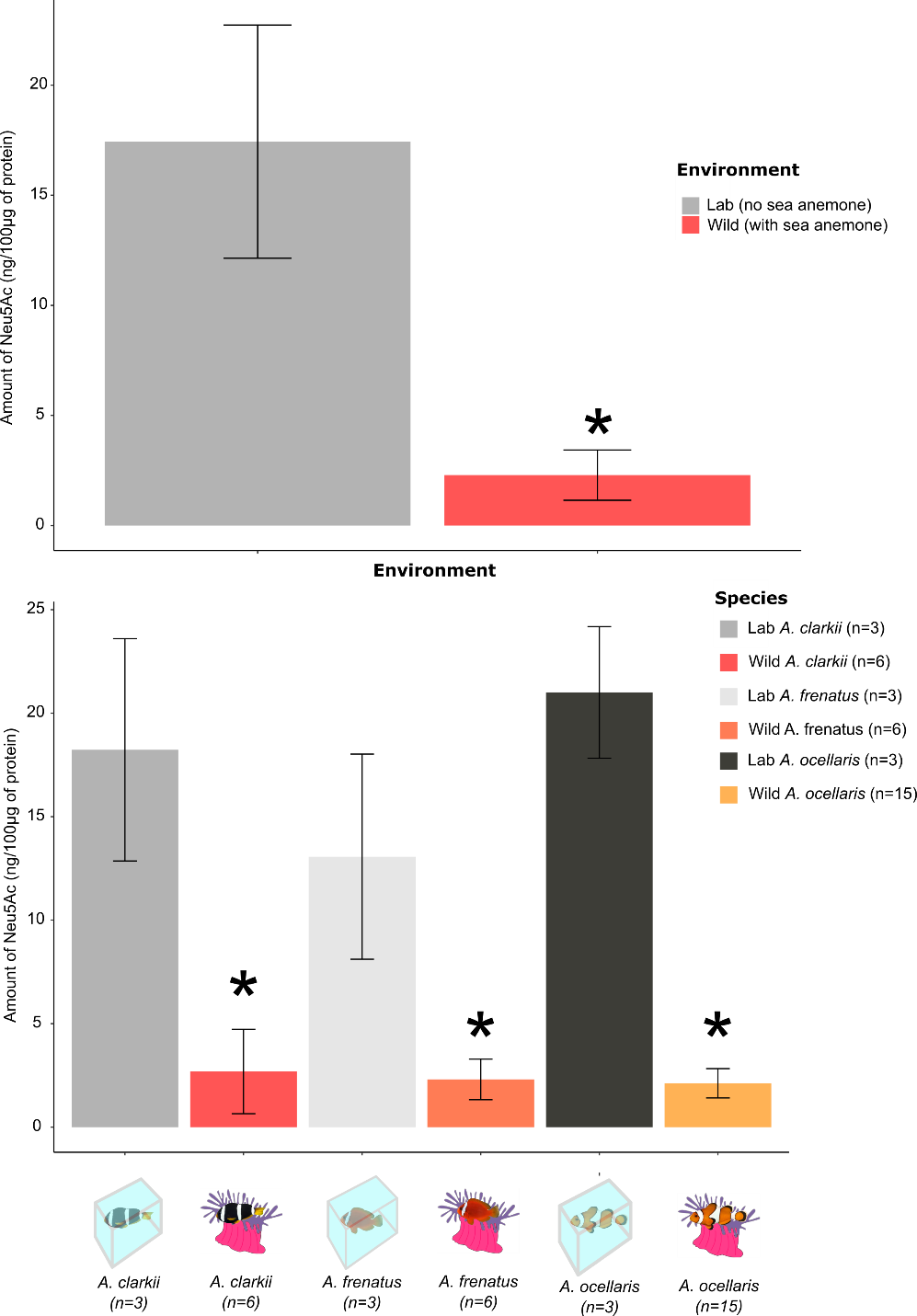

**Supplementary Figure 1:** A) Comparison of Neu5Ac levels between lab maintained anemonefish species and wild caught anemonefish species (*Amphiprion clarkii, Amphiprion frenatus, Amphiprion ocellaris*). A non-parametric Kruskall-Wallis test followed by a pairwise Wilcoxon text were performed and significant differences between lab and wild caught fishes is displayed by a star (*) for each species. B) Comparison of Kdn levels between organs of the anemonefish *A. ocellaris* and the two damselfish *Chrysiptera cyanea* and *Chromis viridis*. No statistical test was performed as Kdn was only detected in *C. viridis*.

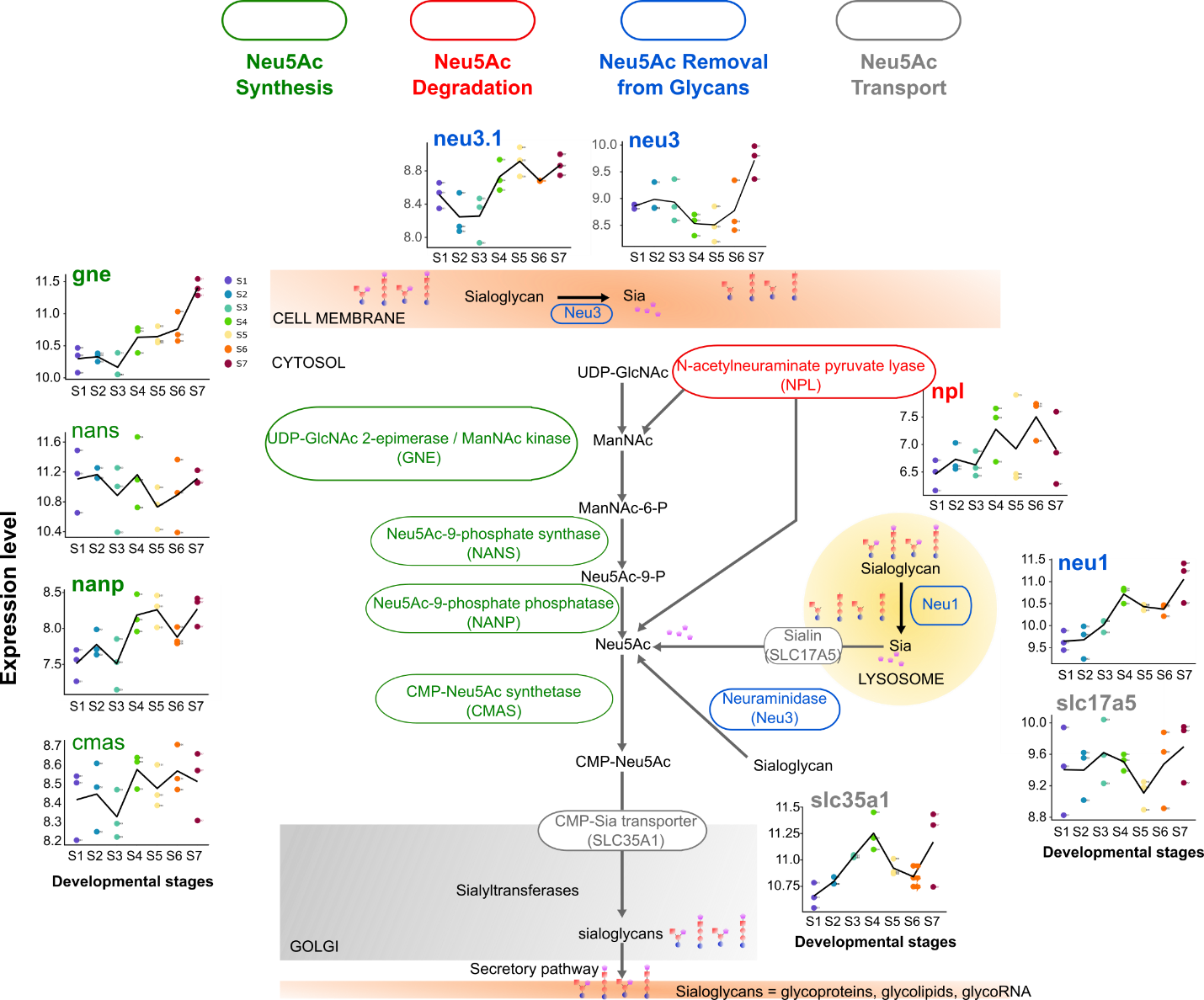
**Supplementary Figure 2:** Complete signaling pathway of Neu5Ac metabolism associated with gene expression levels retrieved from transcriptomic data. Green genes are involved Neu5Ac synthesis, red genes in Neu5Ac degradation, blue genes in Neu5Ac removal from sialoglycans and gray genes in Neu5Ac transport Genes written in bold are significantly differentially expressed between pre metamorphosis stage (S1 and/or S2 and/or S3) and metamorphosis stages (S5 and/or S6 and/or S7).

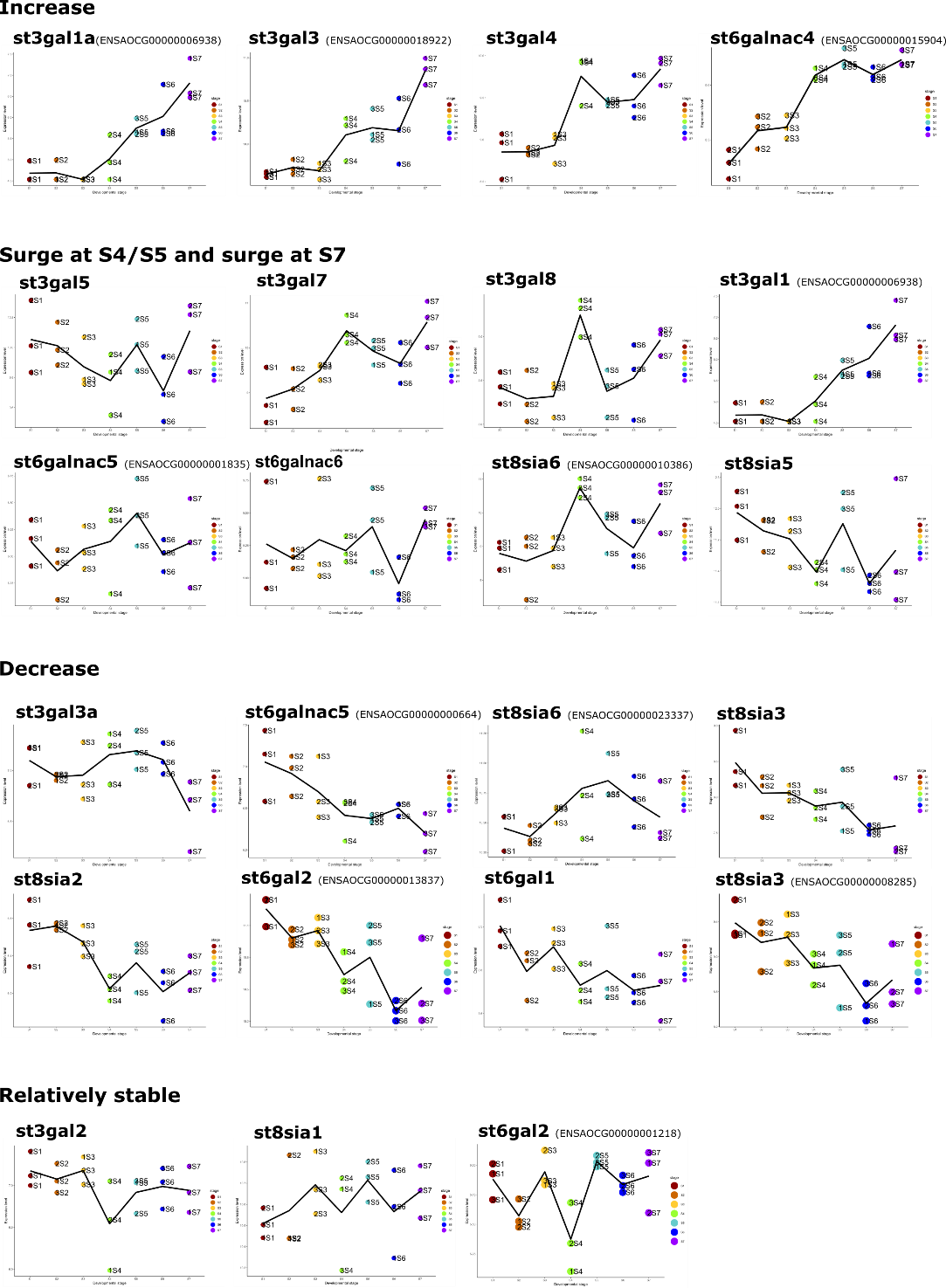

**Supplementary Figure 3:** Expression levels of genes encoding for enzymes, called sialyltransferase, involved in the transfer and fixation of Neu5Ac on sialoglycans. Expression levels were classified into 4 categories (increase, surge at Stage 4 or 5, decrease, and relatively stable). Genes written in bold are significantly differentially expressed between pre metamorphosis stage (S1 and/or S2 and/or S3) and metamorphosis stages (S5 and/or S6 and/or S7).

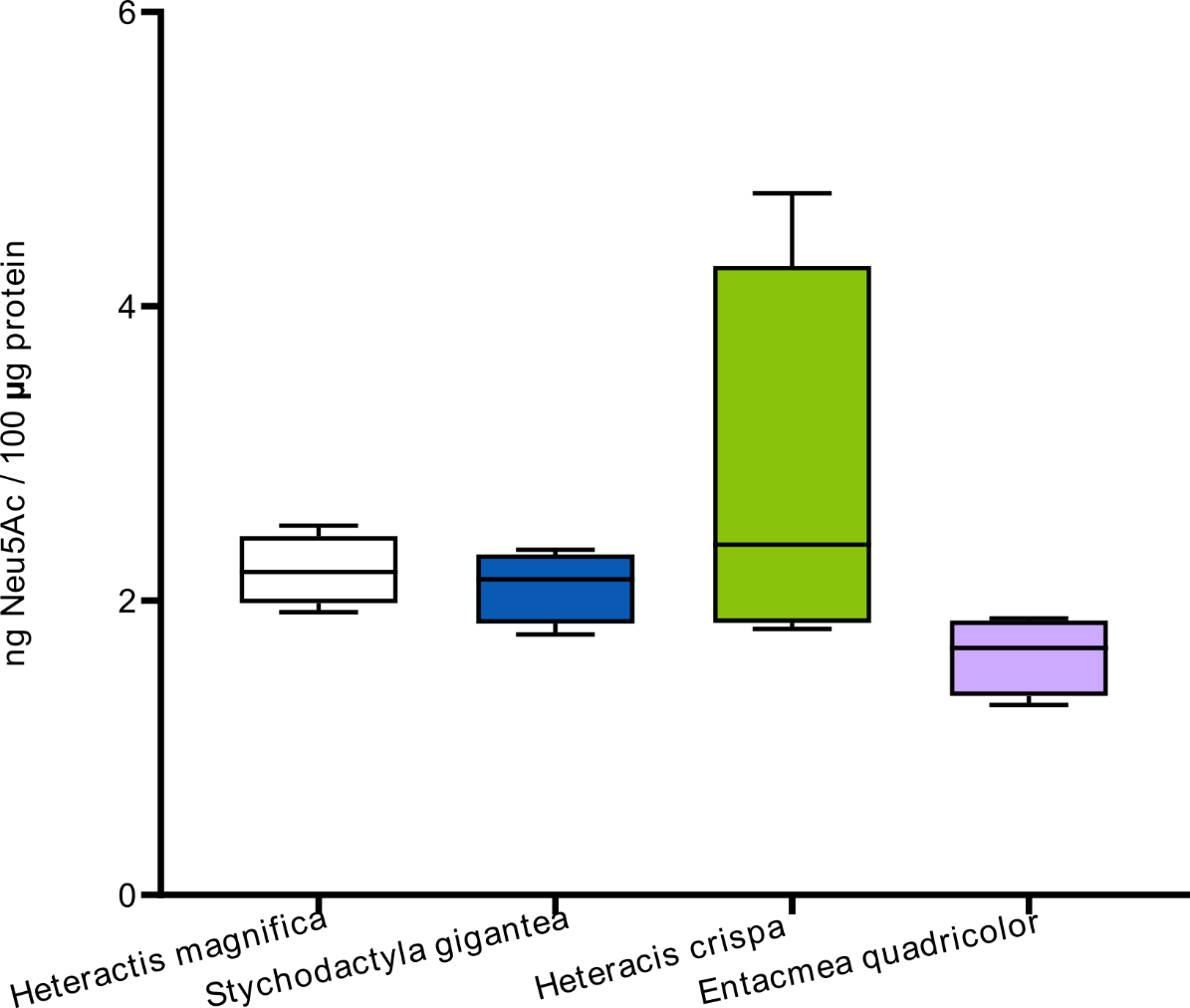

**Supplementary Figure 4: SA levels below detection limit in Giant sea anemones.** Neu5Ac levels in *Heteractis magnifica, Stichodactyla gigantea, Heteractis crispa* and *Entacmea quadricolor.* A non-parametric Kruskall-Wallis test was performed and was not significant (*p-value*=0.1).

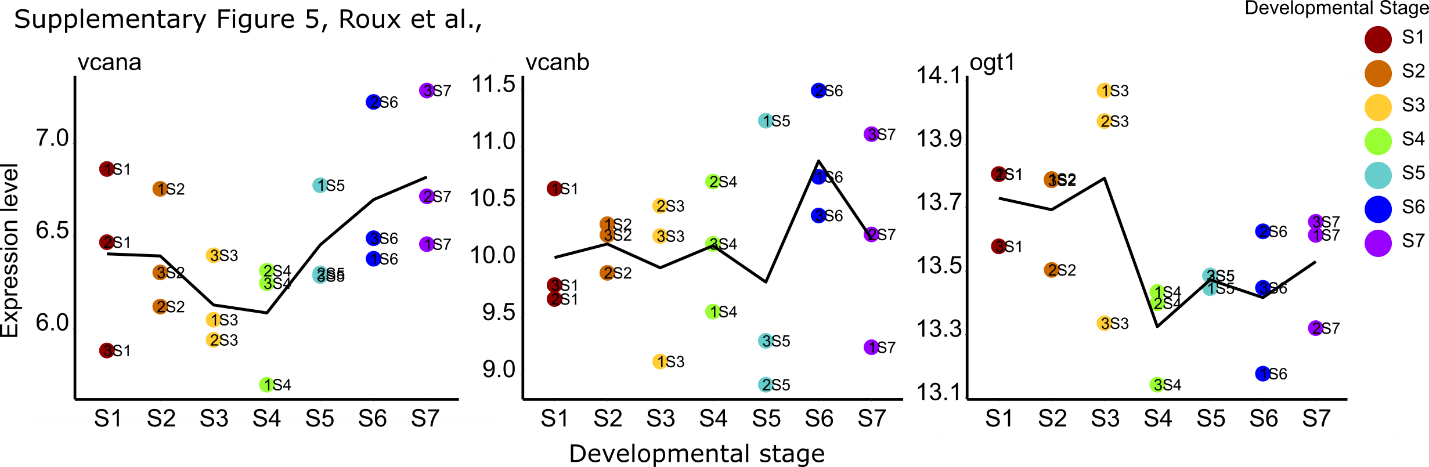

**Supplementary Figure 5:** Expression levels of genes encoding for versican core protein (*vcan*) and the O-GlcNAc transferase (*ogt*). Expression of versican core protein in clownfish skin is thought to bind to N-acetylated sugars and could therefore mask them to the host chemoreceptors therefore preventing nematocyst discharge. The protein O-GlcNAcase has the potential to cleave N-acetylated sugars from different cell surface molecules. Genes written in bold are significantly differentially expressed between pre metamorphosis stage (S1 and/or S2 and/or S3) and metamorphosis stages (S5 and/or S6 and/or S7).
